# Polygamous breeding system identified in the distylous genus *Psychotria*: *P. manillensis* in the Ryukyu archipelago, Japan

**DOI:** 10.1101/2020.10.14.334318

**Authors:** Kenta Watanabe, Akira Shimizu, Takashi Sugawara

**Affiliations:** National Institute of Technology, Okinawa College, 905 Henoko, Nago, Okinawa 905-2192, Japan; Department of Biological Sciences, Graduate School of Science and Engineering, Tokyo Metropolitan University, 1-1 Minami-Ohsawa, Hachioji, Tokyo 192-0397, Japan; Research Institute of Evolutionary Biology, Inc., Setagaya-ku, Tokyo, 158-0098 Japan; University Museum, the University of Tokyo, Bunkyo-ku, Tokyo, 113-0033 Japan; Department of Botany, National Museum of Nature and Science, 4-1-1, Amakubo, Tsukuba, 305-0005 Japan

**Keywords:** breeding system, distyly, monoecism, polygamous, *Psychotria manillensis*, Rubiaceae

## Abstract

Distyly is a genetic polymorphism composed of long- and short-styled flowers in a population. The evolutionary breakdown of distyly has been reported in many taxa, and mainly involves a shift toward monomorphism or dioecism. However, a shift toward monoecism has not been reported in distylous species. *Psychotria* (Rubiaceae), one of the world largest genera, consists of distylous species and their derivatives. In our preliminary study, however, we identified some monoecious individuals in a population of *Psychotria manillensis*. To understand the breeding system and reproductive biology of *P. manillensis*, we investigated floral traits, open fruit set, and flower visitors, and performed hand pollination and bagging experiments in five populations of Okinawa and Iriomote islands, Ryukyu Islands, Japan. The populations of *P. manillensis* were composed mainly of monoecious individuals (54%), followed by female (30%), male (14%), and hermaphroditic (2%) individuals at the time of flower collection. Of the collected flowers, 93% were functionally unisexual (male or female), whereas only 6.5% were perfect (hermaphroditic). However, some individuals changed sex mainly towards increasing femaleness during the flowering period. Moreover, 35% of the studied plants changed their sexual expression over the years. *P. manillensis* showed self-compatibility and no agamospermy. The fruit set under open pollination varied among populations and years (1.8–21.9%), but it was significantly higher than that of auto-selfing (0.68–1.56%). Wasps and flies were the main flower visitors and probably the main pollinators of the species. In conclusion, *P. manillensis* was revealed to be polygamous, involving monoecious, female, male, and hermaphroditic individuals. This is the first report of the polygamous breeding system not only in the genus *Psychotria*, but also in all heterostylous taxa.

## Introduction

Heterostyly is a genetic polymorphism composed of two (distyly) or three (tristyly) distinct floral morphs (Darwin 1877; Barrett 1992; Barrett 2019). Distyly is composed of a “long-styled (pin) morph” with the stigma above the anthers and a “short-styled (thrum) morph” with anthers above the stigma (Lloyd and Webb 1992; Naiki 2012). The heights of the anthers and stigmas are reciprocal between the two morphs, and the species usually have self- and intramorph incompatibilities (Ganders 1979; Barrett et al. 2000; Barrett et al. 2002). These features are a mechanism for promoting outbreeding and avoiding self-fertilization (Barrett 1992), and distyly has been reported for more than 28 flowering plant families (Barrett et al. 2000; Naiki 2012). The genus *Psychotria* L. (Rubiaceae) is one of the largest distylous genera in the world, with more than 1,800 species (Davis et al. 2001 2009; Nepokroeff et al. 2003; Razafimandimbison et al. 2014), most of which are either distylous or have breeding systems derived from distyly (Hamilton 1990; Watanabe and Sugawara 2015).

The evolutionary breakdown or modification of distyly has occurred in many taxa, and several different patterns have been documented (Ganders 1979; Naiki 2012). Of these, monomorphism (homostyly or long-/short-styled monomorphy), which is the most frequently reported evolutionary transition (Vuilleumier 1967; Naiki and Nagamasu 2004; Nakamura et al. 2007; Naiki 2012), has been identified in *Psychotria* (Hamilton 1990; Sakai and Wright 2008; Consolaro et al. 2011). The evolution of dioecy from distyly has also been reported for several lineages (Watanabe and Sugawara 2015), including the Hawaiian *Psychotria* (Beach and Bawa 1980; Muenchow and Grebus 1989; Sakai et al. 1995; Wagner et al. 1999) and *P. asiatica* in Japan (Watanabe et al. 2014b). However, there is no report of the monoecism, not only in the genus *Psychotria* but also in any distylous taxon in the flowering plants.

Our preliminary observations revealed that several individuals of *P. manillensis* have both male and female flowers in a single inflorescence, and consequently, this species may be monoecious. Although most individuals of *P. manillensis* have male- or female-like flowers, some have perfect flowers. Additionally, the sexual forms are often variable among the individuals. Since the sexual forms of each flower and/or individual in *P. manillensis* are irregular and complex, more research is required to clarify the breeding system and reproductive mechanism.

There are four *Psychotria* species other than *P. manillensis* in Japanese subtropical islands: *P. homalosperma*, (Watanabe et al. 2014a; Watanabe et al. 2018), *P. boninensis* (Kondo et al. 2007; Sugawara et al. 2014), and *P. serpens* (Sugawara et al. 2013, 2016) are both morphologically and functionally distylous; *P. asiatica* (Watanabe et al. 2014b; the latter species in Japan, Taiwan and Vietnam had been recognized as *P. rubra*, but now it is regarded as a synonym of *P. asiatica*) is functionally dioecious with distylous morphology. DNA sequencing showed that *P. manillensis* is most closely related to dioecious *P. asiatica*, although the detailed phylogenetic relationship of the two species remains unclear due to lack of data from the Philippines (Watanabe et al. unpublished data). *P. asiatica* in the Ryukyu Islands is tetraploid (2n = 42), whereas *P. manillensis* is octaploid (2n = 84) and F_1_ interspecific hybrids between the two species (hexaploid, 2n = 63) have been also reported in the Ryukyu Islands (Nakamura et al. 2003). Although geographical distributions of these two species overlap only in the Ryukyu Islands, the breeding system of *P. manillensis* might be related to the ancestors of *P. asiatica*.

The objectives of this study were to investigate the breeding system and the reproductive mechanism of *P. manillensis* in natural populations of the Ryukyu Islands, Japan. Does this species have the polygamous breeding system with monoecious individuals? How does this species reproduce in the field? To answer these questions, our study tested the following predictions:: 1) stigmas with poorly developed stigmatic papillae have no female function, 2) flowers can be classified into four sexual types (male, female, perfect and sterile) based on combinations of stigma and anther types, 3) *P. manillensis* has the polygamous breeding system consisting of monoecious, female, male and hermaphroditic plants, 4) *P. manillensis* has self-compatibility but not autogamy, 5) the insect pollination increases fruit set in the field.

## Materials & Methods

### Study plants and sites

*P. manillensis* (Bartl. ex DC.) is a woody shrub, 2–3 m in height, commonly found in limestone evergreen forests with alkali soil and widely distributed from the Ryukyu Islands to the Lanyu and Ludao Islands of Taiwan and the Philippines (Yamazaki 1993). Inflorescences are terminal and each possesses 20–150 flowers (Fig. 1). Corollas are white to white-green and have a short funnel shape, 2–3 mm in length, with white hairs inside. Flowers bloom for a single day; they open in the morning and fade in the evening. Normally, one to five flowers in a single inflorescence bloom each day, and one inflorescence blooms for about two weeks. The flowering season is from June to August, and fruits mature from November to January on Okinawa Island.

**FIGURE 1.**
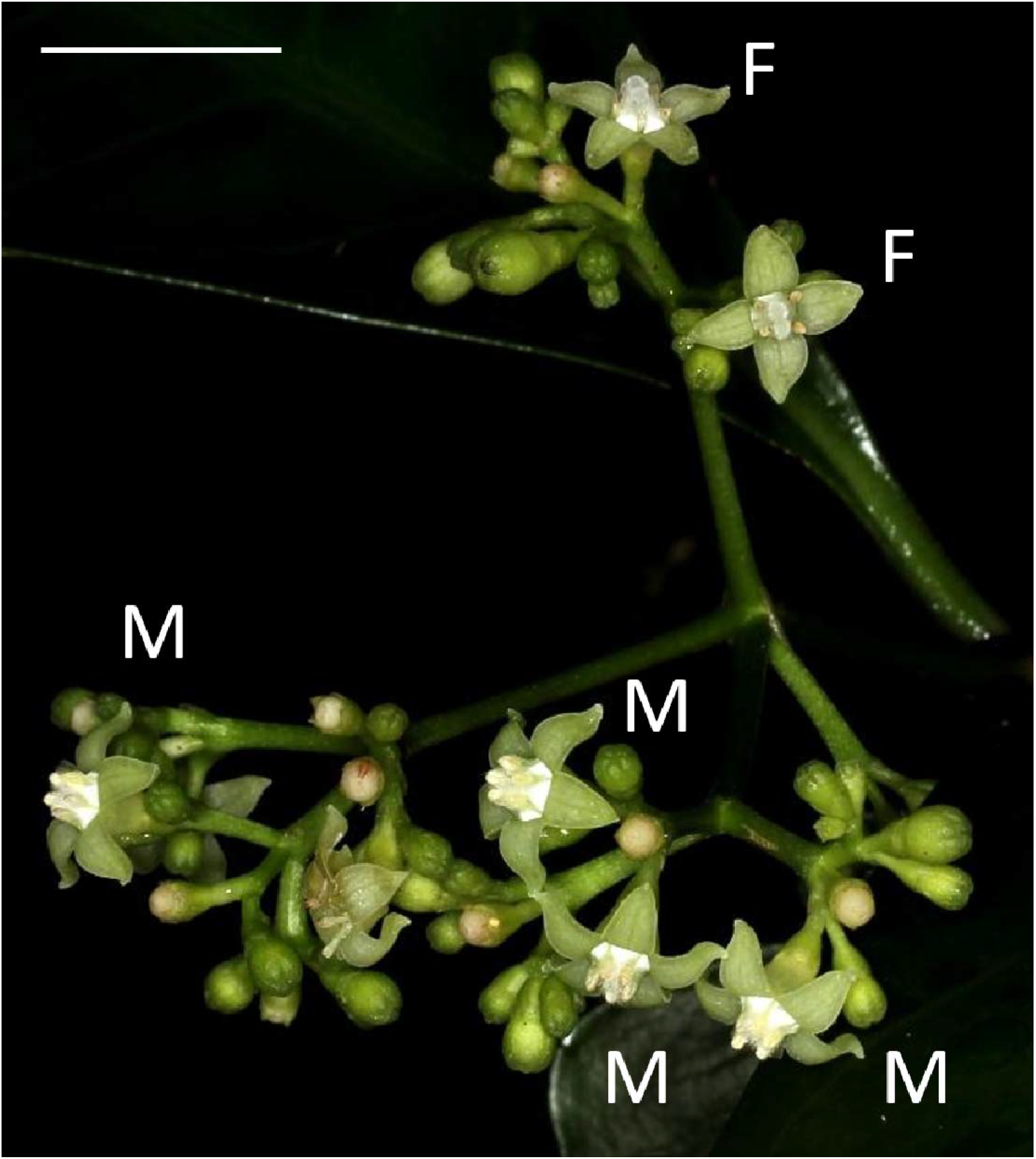
Flowers of *Psychotria manillensis* on Iriomote Island, Japan. Male-like flowers (M) with poorly developed stigmas and anthers with pollen sacs full of pollen grains and female-like flowers (F) with well-developed stigmas and anthers with no pollen grain in a single inflorescence. Scale bar = 10mm.

We examined three populations on Okinawa Island: the Katsuu population at Mt. Katsuu-dake (26.63°N, 127.94°E, 310 m above sea level [a.s.l.]), the Oppa population at Mt. Oppa-dake (26.67°N, 127°96′E, 250 m a.s.l.) and the Sueyoshi population at Sueyoshi Park (26.23°N, 127.72°E, 65 m a.s.l.), and two populations located on Iriomote Island: the Sonai population at the Sonai beach forest (24.37°N, 123.75′E, 5 m a.s.l.) and the Uehara population at the Uehara forest (24.39°N, 123.81°E, 40 m a.s.l.). These three populations on Okianwa Island and two populations on Iriomote Island are representative of limestone forest of those islands, and populations are well separated by other types of forest. All field studies were performed from 2011–2013. Field experiments were approved by Okinawa Prefecture (permit number 23) and Naha city (permit number 272).

### Classification of stigmas and anthers of flowers

We collected 294 flowers from 20 individuals (14 or 15 flowers per individual) in the Sueyoshi population in June–August 2011. We observed the sexual organs of all the collected flowers under a light microscope (SMZ800; Nikon, Tokyo, Japan) and the unfixed stigmas under a scanning electron microscope (SEM, S-3000N; Hitachi, Tokyo, Japan) to examine their surface structure.

We classified the stigmas of *P. manillensis* into three types based on the degree of development of the stigmatic papillae: I, well-developed; II, moderately developed; and III, poorly developed (Fig. 2). We hypothesized the type III stigma does not have female function. We also classified the anthers of *P. manillensis* into three types based on the size of pollen sacs and the amount of pollen: a, anthers with large pollen sacs (usually more than 10 mm in length) full of pollen grains; b, anthers with small pollen sacs (usually less than 5 mm in length) with half or less of the total capacity of pollen grains; and c, anthers with no pollen sac nor grains (without male function) (Fig. 3). Thus, there were nine possible combinations of stigma and anther types; Ia, Ib, Ic, IIa, IIb, IIc, IIIa, IIIb, IIIc (Fig. S1).

**FIGURE 2.**
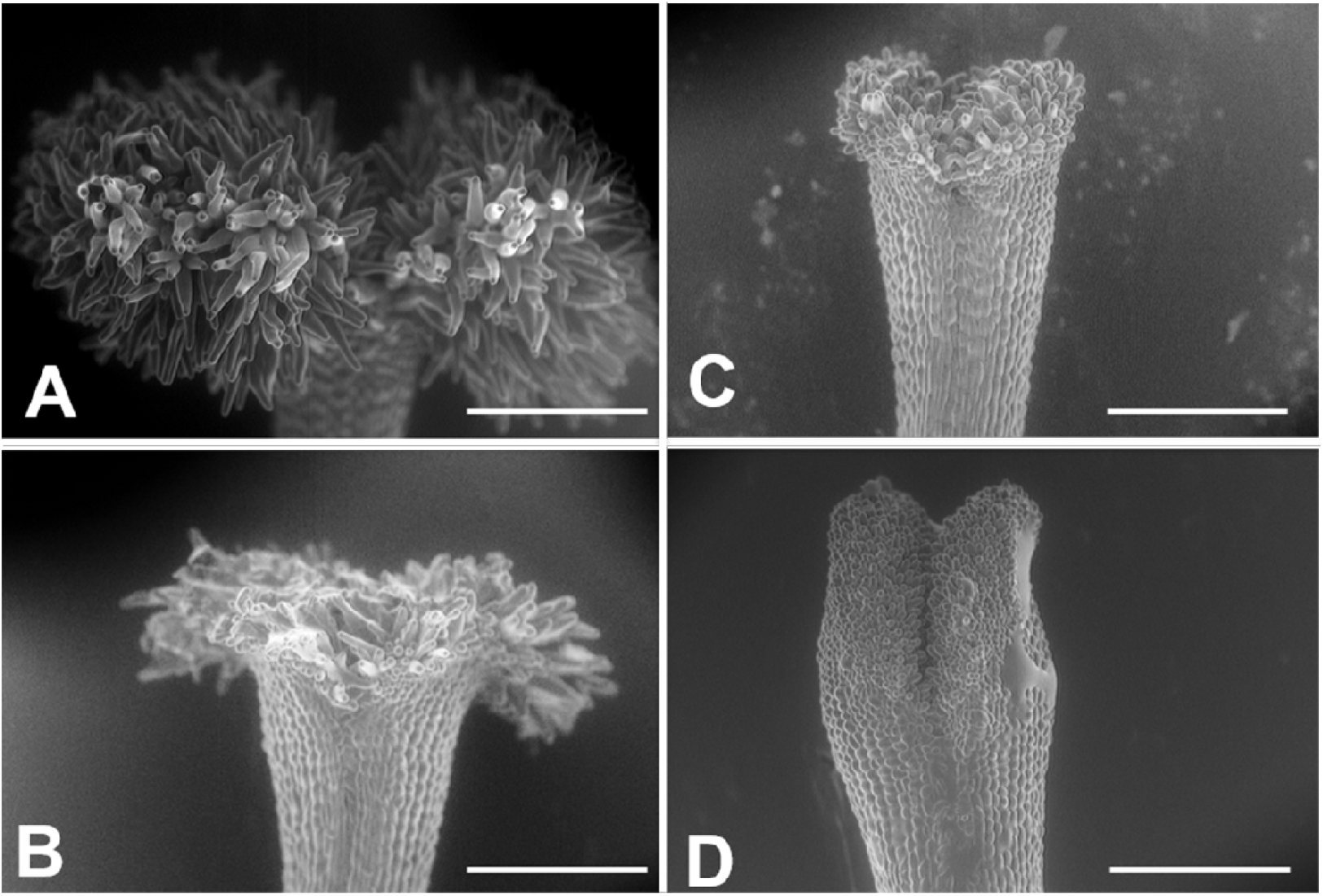
Three types of stigmas in *Psychotria manillensis*. Scanning electron microscopic images of stigmas: A, type-I stigma with well-developed stigmatic papillae; B and C, type-II stigma with moderately developed stigmatic papillae; D, type-III stigma with poorly developed stigmatic papillae. Scale bar = 500 μm.

**FIGURE 3.**
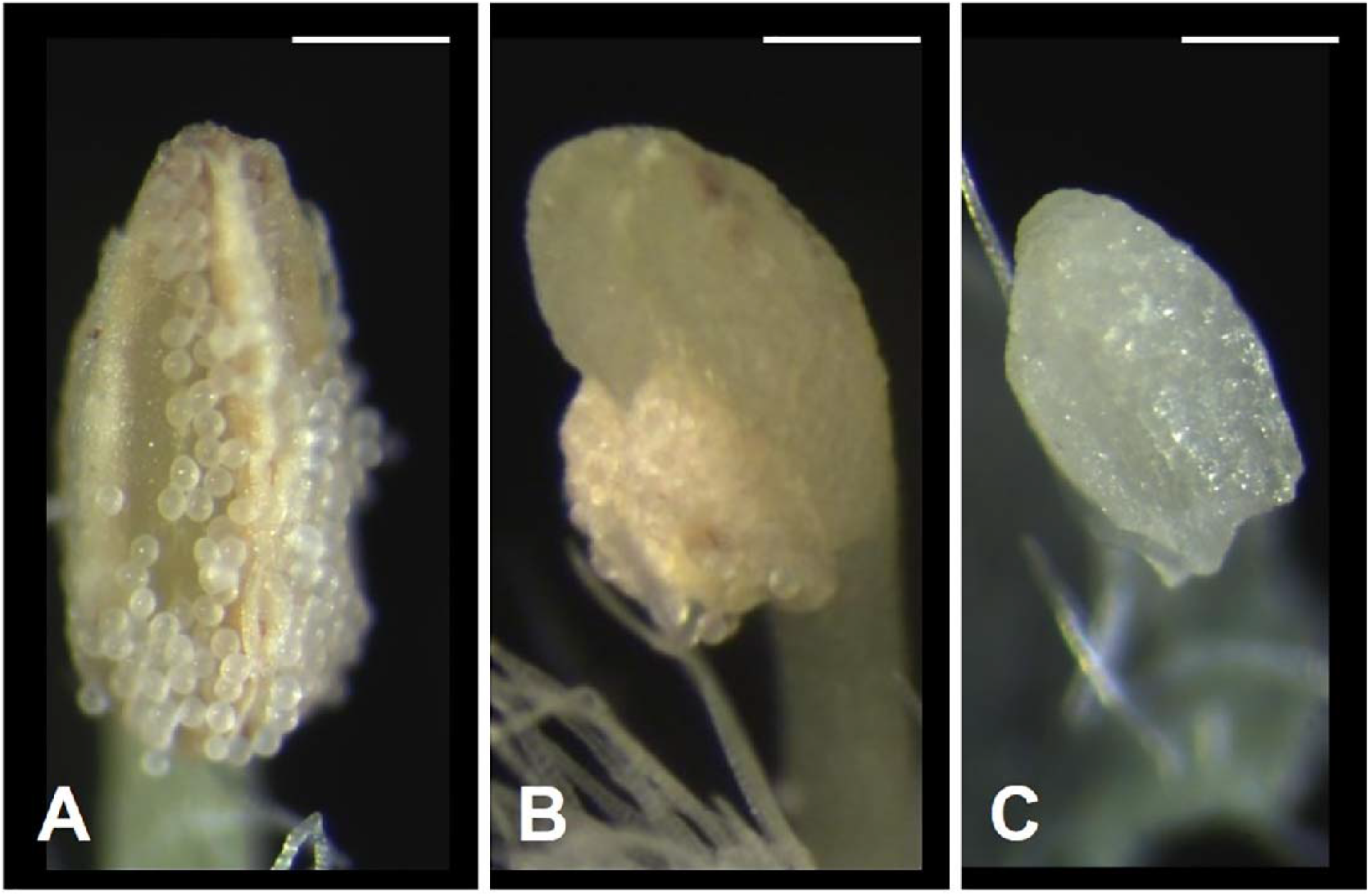
Three types of anthers in *Psychotria manillensis*. A, type-a anther with pollen sacs full of pollen grains; B, type-b anther with small pollen sacs half filled or less with pollen grains; and C, type-c anther with no pollen sac nor grain. Scale bar = 0.3 mm.

### Function of stigmas

To evaluate the function of each stigma type, we cross-pollinated 103 flowers with type-I stigmas (N: Number of individuals = 21), 111 flowers with type-II stigmas (N = 20), and 96 flowers with type-III stigmas (N = 21) at the Oppa and Katsuu populations by transferring pollen from other individuals. Because the stigmas of some flowers were hidden within the corolla tubes, we partially cut the floral tube for pollination. Approximately 50 d after pollination, we counted the number of immature fruits to evaluate the fruit set percentage.

To determine the ability of pollen tubes to access the styles and ovaries, 22 flowers with type-I stigma (N = 10), 17 flowers with type-II stigma (N = 10), and 18 flowers with type-III stigma (N = 10) were cross-pollinated in the laboratory. After 24 hours, the materials were fixed in 5: 5: 90 formalin: glacial acetic acid: 70% ethanol, and then transferred to 70% ethanol for storage. The fixed styles were softened in 8 N NaOH at 4°C for 24 h, washed with water, transferred to 0.005% aniline blue in Na_2_HPO_4_ (pH 11), and stored at 4°C for at least 24 h. The styles were mounted on a slide in glycerol beneath a cover glass and gently pressed to spread the tissue. Pollen grains on the stigmas and pollen tubes in the styles were observed under a fluorescence microscope (Axio Imager Z2; Zeiss, Oberkochen, Germany). Then, the numbers of pollen tubes which reached at the base of the style were counted.

### Flower collection and counts of flowers by type

We collected flowers from 20 previously marked individuals from each of the five populations in 2012 and from the three populations on Okinawa Island in 2013. We visited each plant at least five times (days), each covering the flowering period (early, mid and late), and collected all available flowers (50-396 flowers each). We classified them into the nine flower types as defined above (see Fig. S1). Then, we classified the individuals by gender based on the flower sexual type(s) at the time of collection: female individuals (only female flowers), male individuals (only male flowers), monoecious individuals (both male and female flowers and often with some perfect flowers), and hermaphroditic individuals (only perfect flowers). To calculate the relative femaleness (G) of each plant, we used proportions of female, male and perfect flowers in total number of flowers. Although these proportions cannot represent the “functional femaleness” of the individuals exactly (Lloyd 1980), they can be the indicator of relative femaleness. We used the following formula to calculate the relative femaleness (G): G = (F + P/2) / (F + M + P), where F, P and M are the number of female, perfect and male flowers per plant, respectively. Since the perfect flowers have both female and male function, we assumed the perfect flower has the half of the femaleness of the female flower..

### Measurements of floral traits

Male and female flowers of dioecious species that were derived from distylous ancestors often still have reciprocal positioning of the anther and stigma height. To understand the relationship between morphological traits (anther and stigma heights) and sexual types of flowers (to check for remnants of distylous traits in this species), we measured the anther and stigma height of 12-15 flowers from six individuals (78 flowers in total) of different genders (one female, one male, one hermaphroditic and three monoecious plants) in the Sueyoshi population.

### Self-compatibility and agamospermy test

To test self-compatibility, we self-pollinated 136 flowers (34 Ic, 63 IIc, and 31 IIIa) from 31 individuals in the Oppa population. Inflorescences were bagged 1 d before pollination, and open flowers in bags were pollinated with pollen grains from other flowers in a bagged inflorescence of the same plant. Then, the self-pollinated flowers were bagged again, and the fruit set was observed approximately 30 d after pollination. To test agamospermy (apomixis), we emasculated 10 flowers from nine individuals (90 flowers in total) before anthesis and observed the fruit set approximately 50 d after the treatment.

### Fruit set under open pollination and bagging experiments

Fruit set under open pollination was surveyed for three to five inflorescences of 9–31 individuals (average of 19.8 individuals per site per year) from the three populations on Okinawa Island in 2011 and all five populations on Okinawa and Iriomote Islands in 2012–2013. We counted the number of flower buds in inflorescences and the number of immature fruits on the marked inflorescences approximately 50 d after the flowering peak.

A bagging experiment was carried out to test for spontaneous self-pollination/fertilization in 2012. One inflorescence each of 10, 11, and 18 individuals from the Sueyoshi, Katsuu, and Oppa populations, respectively, was bagged using a mesh bag before blooming, and the fruit set ratio (proportion of flowers that produced fruit) was calculated after all flowers faded (approximately 60-70 d after the flowering peak).

### Flower phenology

We marked 13 inflorescences on 10 individuals in the Oppa population (130 inflorescences in total) and recorded the number and sexual type of open flowers every 2 d from June 26 to August 3, 2012. We divided the flowering period into four spans and calculated the relative femaleness (G) as described above for each span and plant.

### Gender shift over the years

We tagged 21, 18, and 19 plants from Katsuu, Oppa and Sueyoshi populations on Okinawa Island respectively for multiple year tracking of the relative femaleness (G) and the gender of each individual.

### Flower visitors

Flower visitors were observed in all five populations over 8 h (from the time flowers opened until they closed; usually between 0700 and 1800), and some individuals were collected for identification. Observations were performed on July 17 and 18, 2011 and on July 14 and 20, 2012 in the Katsuu population; from July 6 to September 18, 2011 and on July 14, 2012 in the Oppa population; on July 9 and 27, 2011, on June 19 and July 1, 2012 in the Sueyoshi population; and on June 2, 3, 15, 16, and 17, 2011 and on June 13, 2013 in the Sonai and Uehara populations.

### Statistical Analysis

We used R v. 4.1.0 (R Core Team, 2021) for all statistical analysis. The effects of stigma types and pollination treatments to fruit set were tested using the generalized linear mixed models (GLMMs, fitting data to a binomial error distribution with log-link function), using the ‘glmer’ function of the R package ‘lem4’. To test the effect of stigma type to fruit set, the GLMMs included stigma type (I, II, and III) as fixed explanatory variables, fruit set as response variable and individual plants as random effect. To test the effects of pollination treatment to fruit set, GLMMs included pollination treatment (cross and self-) as fixed explanatory variables, fruit set as response variable and individual plants and stigma type as random effects. The goodness of fit to models was assessed by the maximum likelihood-ratio chi-square test, using ‘anova’ function of the R package ‘lme4’. To evaluate the effect of each stigma type, we performed a Tukey post hoc test with the ‘glht’ function of the package ‘multcomp’.

We also analyzed the factors that affect fruit set in the field using GLMM (binomial error distribution with log-link function). For the open pollination, the models included the fruit set as the response variable, year and population as the explanatory variables and individual plant as a random factor. We compared four models with different combinations of the explanatory variables: model 1 (population), model 2 (year), model 3 (population and year), and model 0 (null). The model fit was evaluated with the Akaike information criterion (AIC), and the model with the smallest AIC was regarded as the best fit for our data. Then, we tested the effect of the bagging experiment against the open pollination. In this case, the models included the fruit set as the responsible variable, treatment (open / bagging) as the explanatory variables and individual plant and population as nested random factors. Subsequently, we performed the maximum likelihood-ratio chi-square test.

We performed a one-sided Pearson’s correlation test using ‘cor.test’ function of R to evaluate the association between femaleness (G) and fruit set under open pollination in each population. We also performed a two-sided Wilcoxon test (using ‘wilcox.test’ function in R) to test the difference in relative femaleness of individual between years 2012 and 2013.

## Results

### Function of stigmas and anthers by type

After cross-pollination, fruit set was observed in 64.1% and 65.8% of flowers with type-I and type-II stigmas respectively, but only 2.1% of flowers with type-III stigmas (Table 1). Similarly, pollen tubes reached at the base of styles in 100% (22/22) of type-I stigmas and 82% (14/17) of type-II stigmas, but no pollen tube growth was observed in flowers with type-III stigma (0/18).

**Table 1.**
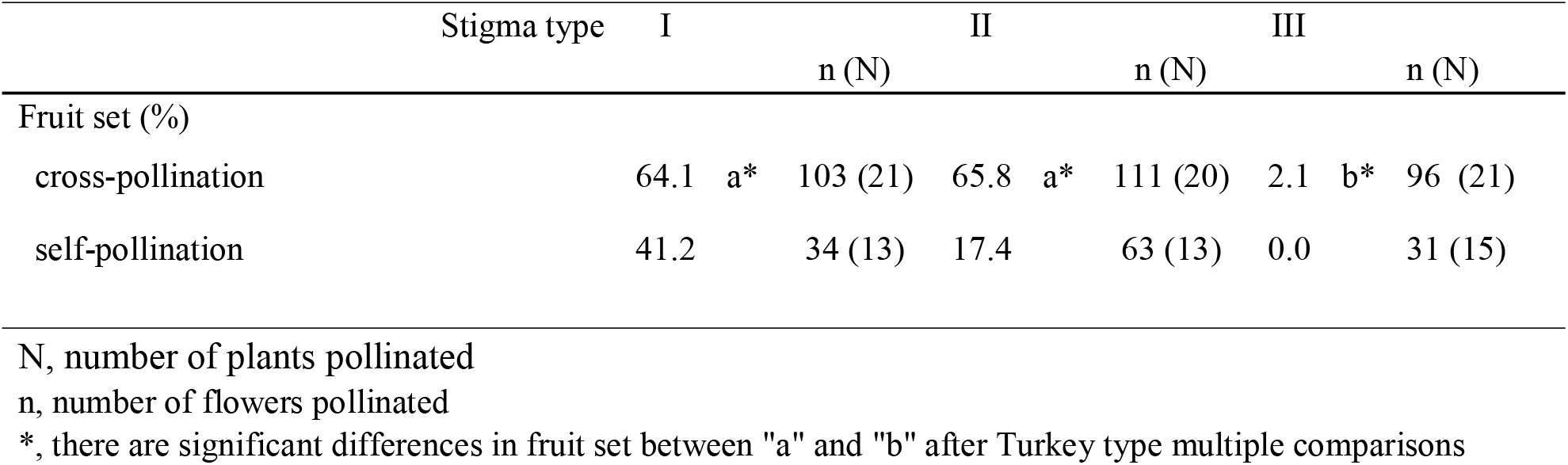
Fruit set after cross- and self-pollination for each stigma type of *Psychotria manillensis* on Okinawa Island, Japan. Type I and II stigmas set more fruits than Type III stigmas after cross-pollination (*P* < 0.0001, binomial GLMM). Fruit set after cross-pollination was greater than that after self-pollination (*P* < 0.0001, binomial GLMM).

|  | Stigma type | I |  |  | II |  |  | III |  |  |
| --- | --- | --- | --- | --- | --- | --- | --- | --- | --- | --- |
|  |  |  |  | n (N) |  |  | n (N) |  |  | n (N) |
| Fruit set (%) |  |  |  |  |  |  |  |  |  |  |
| cross-pollination |  | 64.1 | a* | 103 (21) | 65.8 | a* | 111 (20) | 2.1 | b* | 96 (21) |
| self-pollination |  | 41.2 |  | 34 (13) | 17.4 |  | 63 (13) | 0.0 |  | 31 (15) |

N, number of plants pollinated
n, number of flowers pollinated
\*, there are significant differences in fruit set between "a" and "b" after Turkey type multiple comparisons

Of the nine possible flower types (based on combinations of stigma and anther types, see Fig. S1), eight flower types (namely Ib, Ic, IIa, IIb, IIc, IIIa, IIIb, and IIIc, but not Ia) were found in the field. All flowers had anthers and a stigma, but more than 93% of the flowers differentiated to either male (IIIa and IIIb, 34%) or female (Ic and IIc, 59%). Only 6.5% of the flowers were perfect (Ib, IIa, and IIb) and only 0.5% were sterile (IIIc); no Ia type flowers were observed (Fig. 4). The ratio of type-I and type-II stigmas was 92:8 for the female flowers and 8:92 for the perfect flowers. The proportion of flower types was relatively consistent in all five populations, although the number of perfect flowers was relatively low in the Uehara population (Fig. 5A).

**FIGURE 4.**
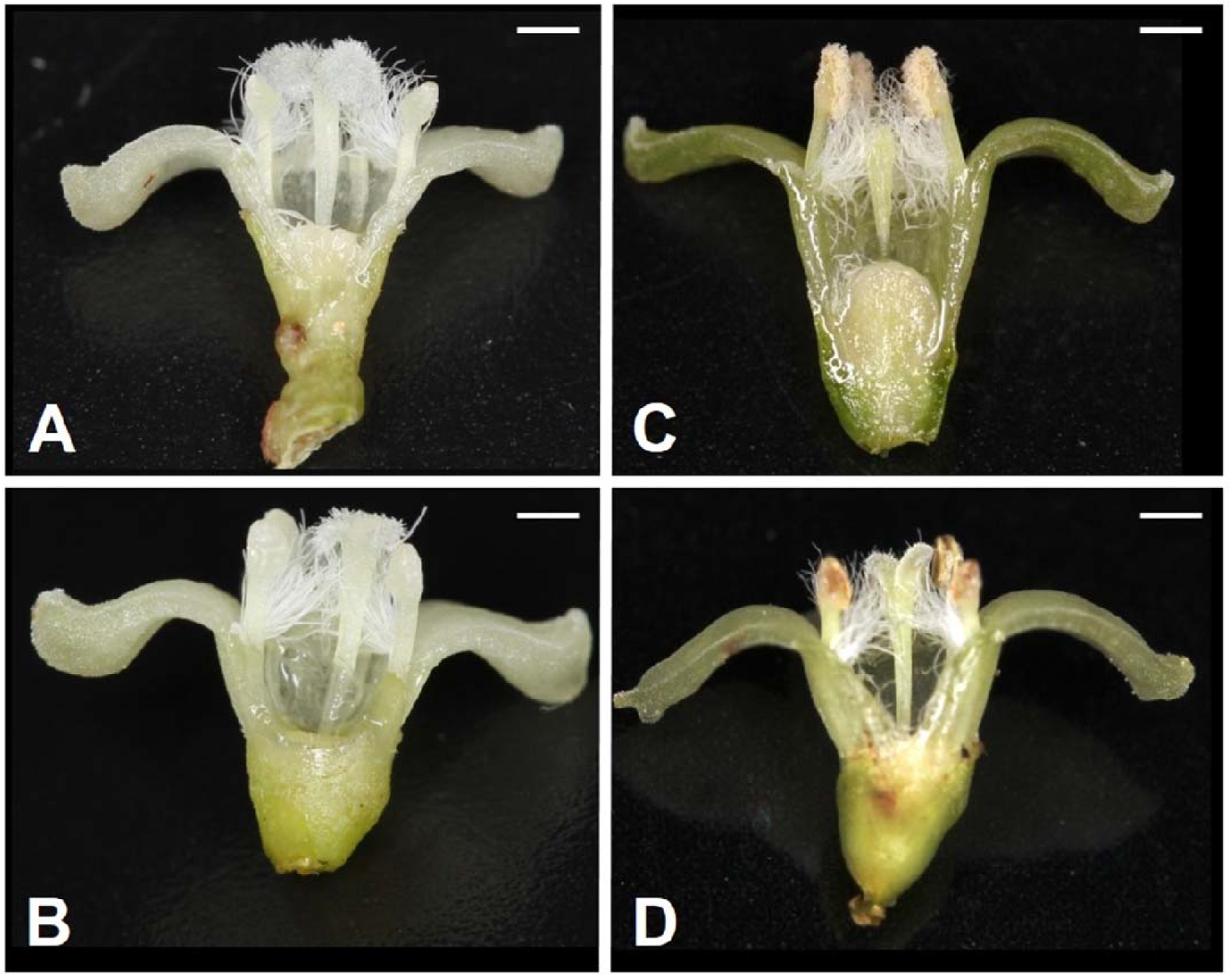
Various flower types in combination with different stigma and anther types of *Psychotria manillensis* on Okinawa and Iriomote Islands, Japan. A, flower with type-I stigma with well-developed stigmatic papillae and type-c anthers with no pollen grain (Ic; 38%); B, flower with type-II stigma with moderately developed stigmatic papillae and type-c anthers with no pollen grain (IIc; 11%); C, flower with type-III stigma with poorly developed stigmatic papillae and type-a anthers with pollen sacs full of pollen grains (IIIa; 32%); D, flower with type-II stigma with moderately developed stigmatic papillae and type-b anthers with small pollen sacs with half or less of the total capacity of pollen grains (IIb; 9%). Ic (A) and IIc (B) flowers function as female, IIIa (C) as male and IIb (D) as hermaphroditic. Scale bar = 1 mm.

**FIGURE 5.**
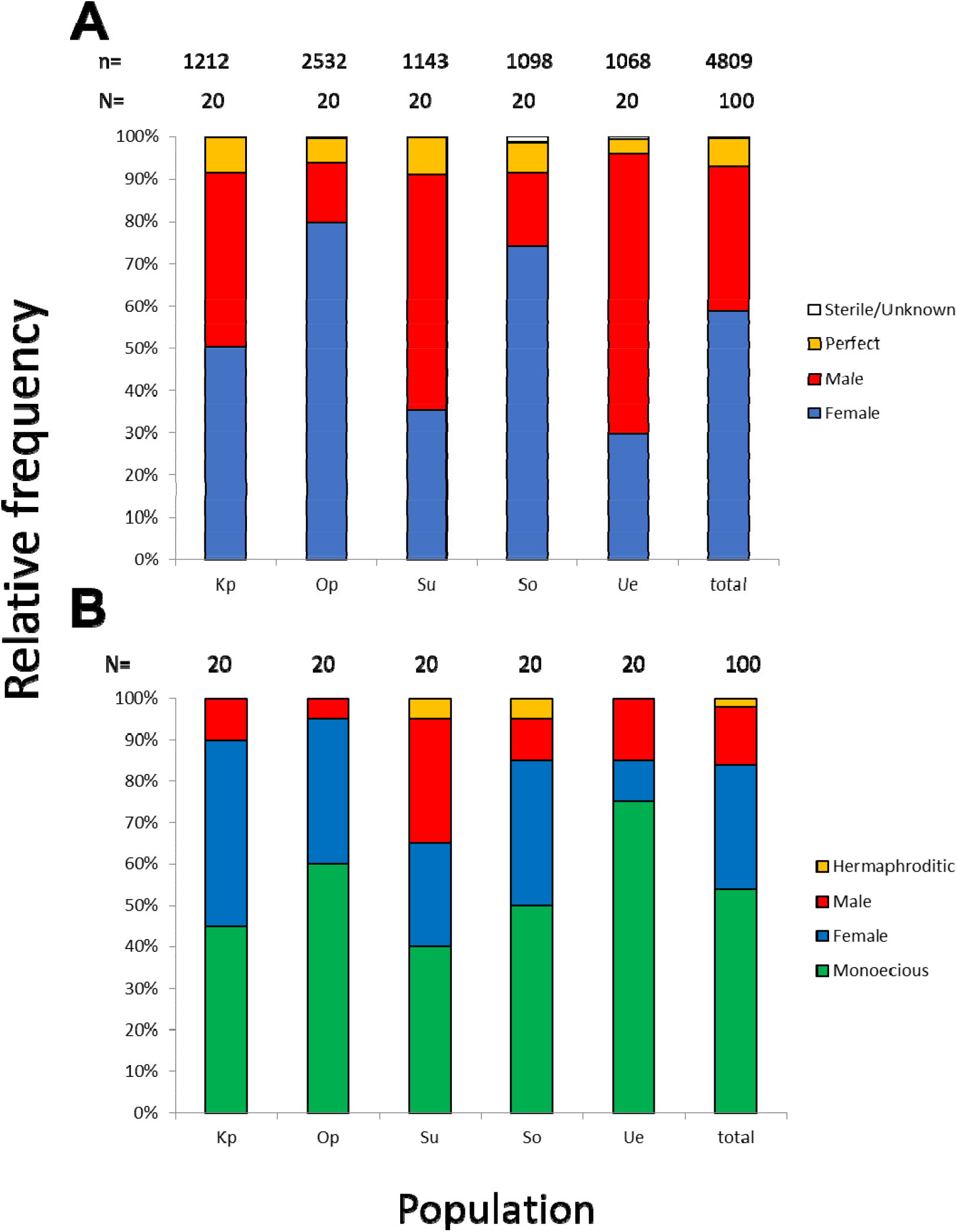
Relative frequencies of female, male, perfect, and sterile flowers (A) and of female, male, monoecious, and hermaphroditic plants (B) in the five populations from Okinawa and Iriomote Islands, Japan. n, number of flowers; N, number of plants. KP, the Katsuu population; OP, the Oppa population; Su, the Sueyoshi population on Okinawa Island. So, the Sonai population; Ue, the Uehara population on Iriomote Island.

Monoecious individuals were the most common (54%) in the three populations, followed by female (30%), male (14%), and hermaphroditic (2%) individuals (Fig. 5B). Female individuals usually consisted of only Ic (female) flowers that had a high stigma with low anthers (Fig. S2A), whereas male individuals consisted of IIIa (male) flowers that had a low stigma and high anthers (Fig. S2B). Of the monoecious individuals, most of the flowers were Ic (female) and IIIa (male), but those types were not distinct when mixed with perfect flowers (Fig. S2C– E).

### Self-pollination and agamospermy test

The fruit set after self-pollination was significantly lower (36.2%) than that after cross-pollination (65.0%; *P* < 0.001, GLMM followed by likelihood ratio test; Table 1). Of 18 studied individuals, 13 showed self-compatibility. Additionally, none of the emasculated flowers set any fruit, revealing that agamospermy did not occur.

### Fruit set under open pollination and bagging experiments

The fruit set under open pollination varied among populations and years (1.8–21.9%; Table S1), and the model 3 which includes both populations and years as explanatory variables showed the smallest AIC (binomial GLMM; Table S2). Some male individuals (only male flowers at the time of flower collection) also produced fruit, although femaleness (G) was significantly correlated with fruit set in all populations (*P* < 0.05, Pearson’s correlation test; Fig. S3), except for the Katsuu population (*P* = 0.35). Some flowers set fruit after bagging experiments, although the fruit set was significantly lower than that under open pollination (*P* < 0.0001, the likelihood ratio test after binomial GLMM; Table S1, S2).

### Flowering phenology and flower types within inflorescence through flowering period

The flowering period of 10 individuals observed in the Oppa population was approximately 42 d (Fig. S4A). Femaleness (G) increased in seven individuals and decreased only in one plant during the flowering period, whereas two individuals produced only female flowers (Fig. S4B). Of 127 inflorescences, 44 (35%) had only female flowers, 44 (35%) had male, female, and perfect flowers, 23 (18%) had male and female flowers, 14 (11%) had male and perfect flowers, and two (1.6%) had only male flowers (Fig. 6).

**Figure 6.**
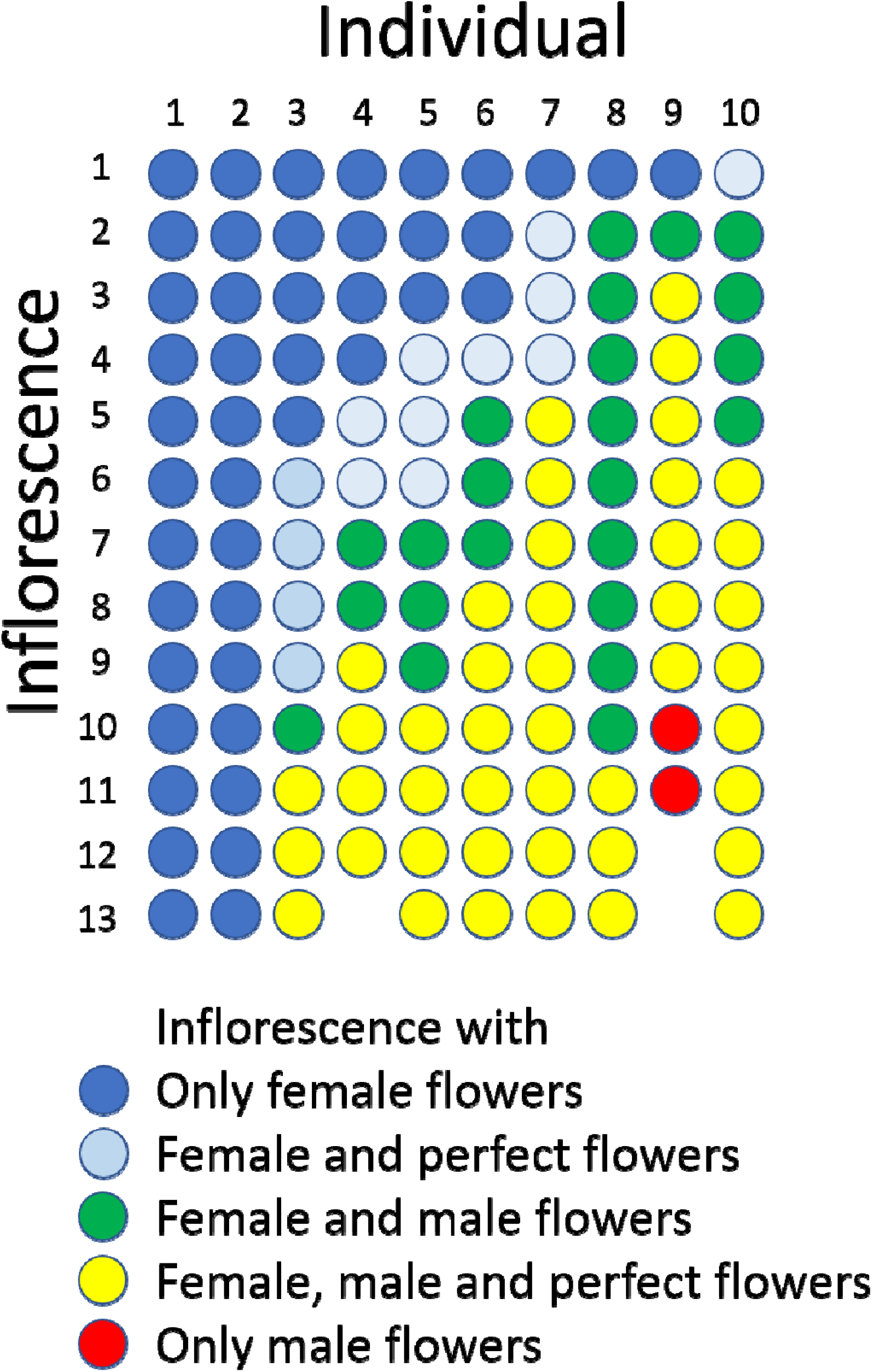
Sexual expression of inflorescences of *Psychotria manillensis* during the flowering period (42 d; June 26–August 3, 2012) in the Oppa population of Okinawa Island, Japan. Ten columns represent ten plants monitored and each circle represents each inflorescence of those plants. The colors of circles indicate the sexual type of inflorescences.

### Gender shift over the years

of the 58 checked plants, relative femaleness increased in 24 plants (41%) and decreased in three plants (5%, Fig. 7A). Although the majority of female and monoecious plants kept the same gender in 2012-2013 season, 20 plants (35%) changed their sex (Fig. 7B). Five female plants changed to monoecious, six monoecious plants to female, one hermaphroditic plant to monoecious, seven male plants to monoecious, and one male plant to female.

**Figure 7.**
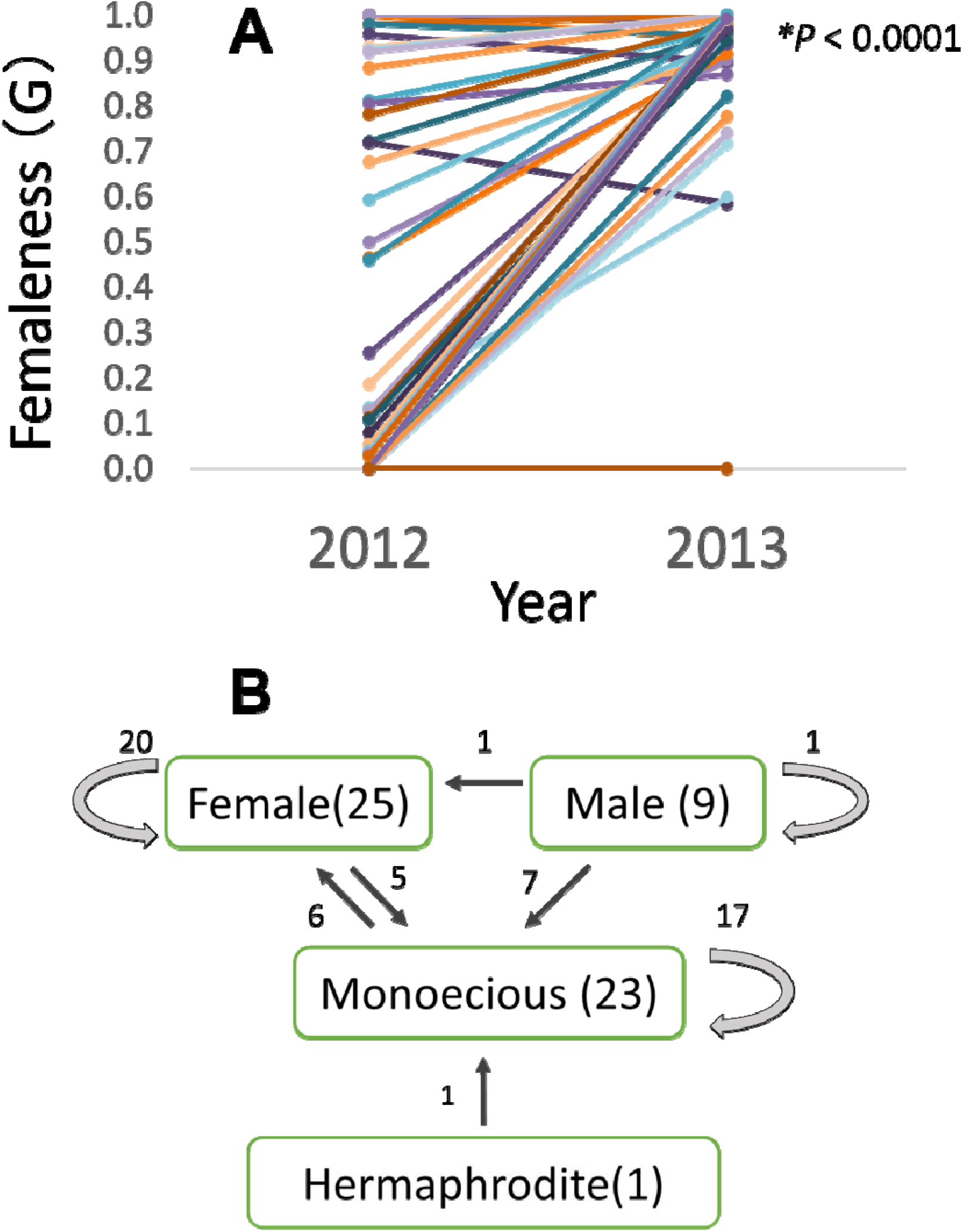
Relative femaleness G (A) and sex changes (B) of 58 plants in three populations on Okinawa Island in 2012-2013. Numbers in parenthesis are the numbers of plants in 2012, numbers with arrows indicate plants changed the sex. Relative femaleness between years was compared using a both sided Wilcoxon test (W=983, *P* < 0.0001).

### Flower visitors

A total of 39 insect species of 20 families belonging to six orders visited the flowers of *P. manillensis* in the five populations (Table S3, Fig. 8). Of these insects, wasps of the family Vespidae (*Vespa* and *Polistes*) and flies of the families Calliphoridae, Syrphidae, and others were the most frequent visitors, and usually pollen grains were attached to their heads. Wasps visited the flowers, sucked nectar, and left quickly for other flowers, whereas flies stayed relatively longer after licking the nectar. Bees (*Apis mellifera, Amegilla* spp., Apidae and *Lasioglossum*, Halictidae) also visited the flowers, but much less frequently than wasps and flies. Bees touched both the stigmas and anthers (usually pollen attached to their heads or bodies) and moved quickly from one flower to another, Butterflies (*Papilio, Lampides*, and *Athymaperius*) and moths (Crambidae) also visited flowers on Okinawa Island, but only occasionally touched the anthers or stigmas with their proboscises. Hawkmoths (*Macrogrossum*), beetles, stink bugs, and crickets were occasional visitors. Only syrphid flies (*Allobaccha nubilipennis* and *Sphaerophoria* sp.) fed on pollen, whereas all others fed on nectar.

**FIGURE 8.**
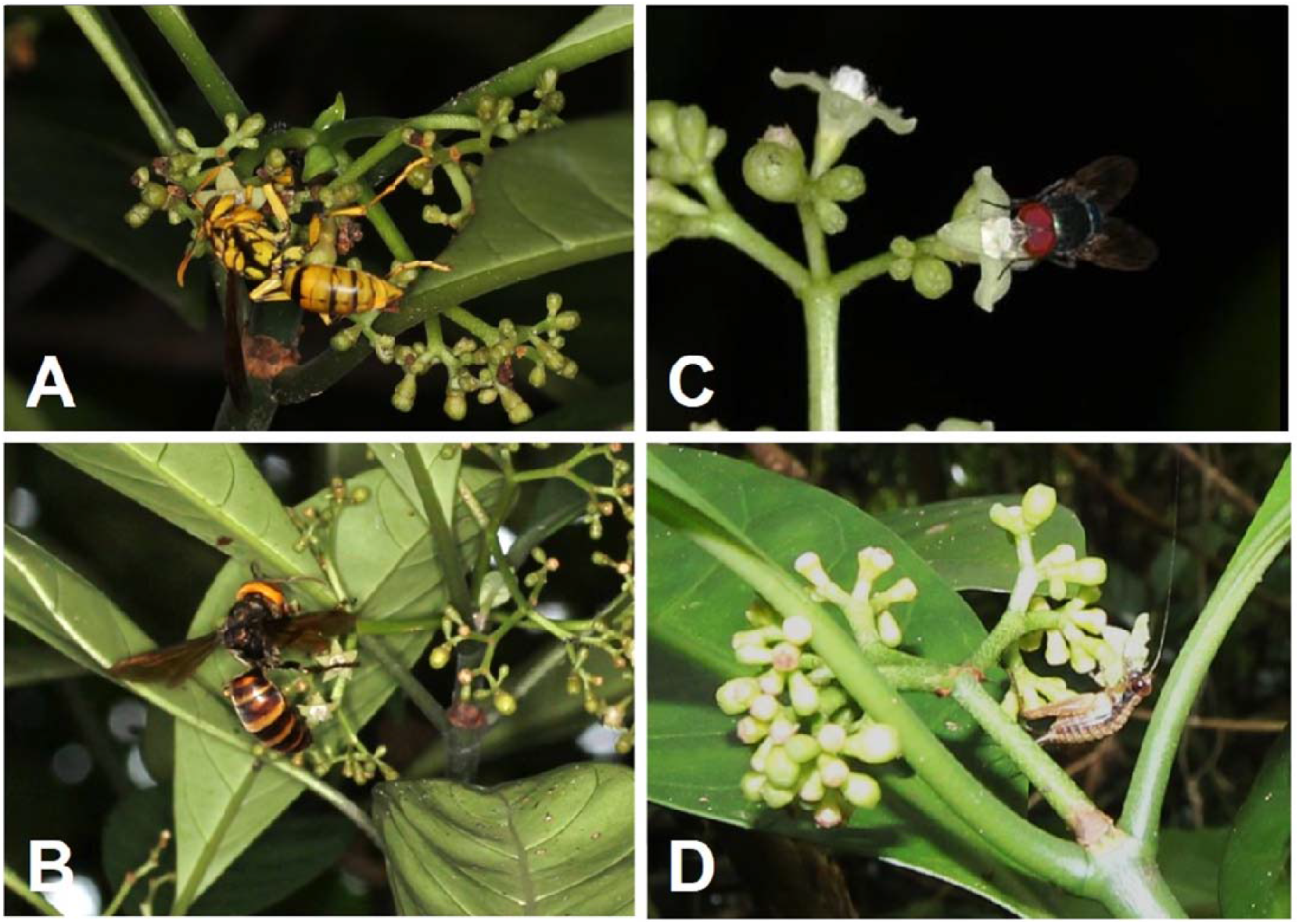
Flower visitors of *Psychotria manillensis* on Okinawa and Iriomote Islands, Japan. A, *Polistes rothneyi* ; B, *Vespa analis* ; C, *Lucilia porphyrina* ; D, *Cardiodactylus guttulus*.

## Discussion

This study aimed to investigate the breeding and floral biology of *P. manillensis* in the natural populations of the Ryukyu Islands. The results showed that *P. manillensis* was self-compatible and polygamous, including monoecious, female, male, and hermaphroditic individuals, and that flowers were visited by wasps and flies. To our knowledge, this is the first report of polygamous plants not only in the genus *Psychotria* but also in all heterostylous taxa worldwide.

The results of cross-pollination indicated that the stigmas of type-I and type-II, but not type-III showed a female function. These results confirmed our hypothesis regarding the distribution of the four sexual flower types; perfect flowers (Ib, IIa, and IIb), male or staminate flowers (IIIa and IIIb), female or pistilate flowers (Ic and IIc), and sterile flowers (IIIc).

Individuals observed in the five populations were monoecious plants with male, female, and often some perfect flowers, female plants with only female flowers, and male plants with only male flowers, whereas hermaphroditic plants with only perfect flowers were found in three populations. Based on these data, the breeding system of *P. manillensis* could be characterized as “polygamous” (Darwin 1877; Sakai and Weller 1999), since no other better terms can be applied to describe the co-existence of monoecious, female, male, and hermaphroditic individuals in a single population (Sakai and Weller 1999).

Self-pollination experiments showed that *P. manillensis* is self-compatible, whereas emasculation experiments indicated that no agamospermy occurs in *P. manillensis*. The fruit set after bagging experiments indicated that auto-self-pollination occurred without any pollinators; however, the fruit set was significantly lower than that under open pollination, revealing that pollinators enhance the reproductive success of *P. manillensis*.

The main flower visitors were wasps (family Vespidae) and flies (family Calliphoridae and others) in all the five populations. Although both wasps and flies licked the nectar, wasps moved more actively among flowers, and probably had a higher contribution to the pollination of *P. manillensis* than flies. Hence, wasps and flies were probably the most important pollinators of *P. manillensis* in the Ryukyu Islands, similar to what occurs in the closely related species *P. asiatica* (Watanabe et al. 2014b). This may cause natural hybridization between these two species in the field.

The adaptive consequences of the polygamous breeding system in *P. manillensis* remain unclear. The imperfect shapes and functions of the reproductive organs indicate the instability and imperfect adaptation of the breeding system. Sex changes in the flowering season and over the years indicate the plasticity of the sexual expression. Although there is no evidence of dichogamy in this species from our data, further investigation about sex change is needed to understand its reproductive role.

Since distyly is the ancestral breeding system in the genus *Psychotria* (Hamilton 1989; Watanabe and Sugawara 2015), it is interesting to know how this polygamous breeding system evolved in *P. manillensis*. It is possible that *P. manillensis* is derived from an autopolyploid *P. asiatica* or an allopolyploid *P. asiatica* and another species (polyploid hybrid). It might be easier to explain the evolution of the polygamous system with partial monoecism in *P. manillensis* from dioecism through polyploidization than directly from distyly. Polyploidization and the evolution of monoecism from dioecism have also been reported to co-occur in *Mercurialis annua* (Pannell et al. 2004). In *M. annua*, the occurrence of male and female flowers within a single plant could be explained by the heterogeneous expression of genes responsible for the sexual determination of flowers that doubled through polyploidization (Russell and Pannell 2015). Theoretically, a similar mechanism may also operate in *P. manillensis*, although further experiments are required to confirm this idea.

Overall, this is the first report revealing the polygamous breeding system in the genus *Psychotria* as well as in all heterostylous taxa worldwide. Understanding the evolution of the breeding system in *P. manillensis* might help to better understand the diversification of the breeding systems derived from distyly. Further research is required to understand the evolutionary pathway, adaptive consequences, and origin of the polygamous breeding system. Future studies should focus on the phylogenetic background and genetic mechanisms for sex determination in *P. manillensis* as well as its relationship with *P. asiatica*.

## Acknowledgements

We thank Dr. S. Shinonaga and Mr. N. Kikuchi for their assistance in insect identification; Dr. A. Iguchi for his advice on statistical analysis; Drs. M. Yokota, T. Denda, K. Nakamura, T. Takaso, and T. Konjo for providing helpful information.

