## Supplemental Figures for "Polygamous breeding system identified in the distylous genus *Psychotria*: *P. manillensis* in the Ryukyu archipelago, Japan"

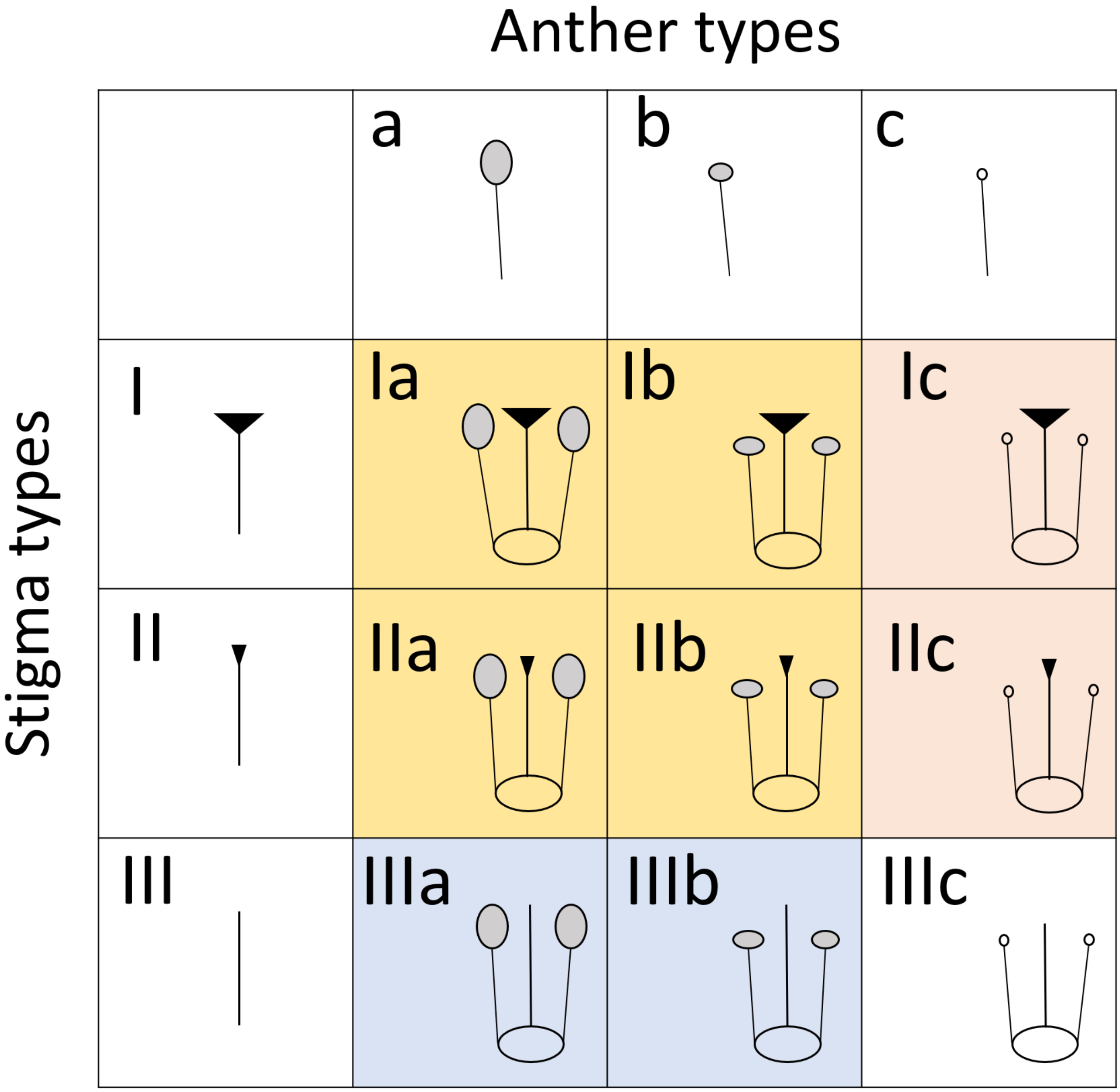


**FIG. S1**

Nine possible flower types with combination of three stigma types and three anther types of *Psychotria manillensis* in the Ryukyu Islands. Three stigma types are: type-I stigma with well-developed stigmatic papillae, type-II stigma with moderately developed stigmatic papillae, type-III stigma with poorly developed stigmatic papillae. Three anther types are: type-a anther with pollen sacs full of pollen grains; type-b anther with small pollen sacs half-filled or less with pollen grains; and type-c anther with no pollen sac nor grain. Supposing that type-III stigmas have no reproductive function, there are four sexual types of flowers: IIIa and IIIb (blue) are functionally male flowers with functional anthers and non-functional stigmas; Ic and IIc (red) are female flowers with non-functional anthers and functional stigmas; Ia, Ib, IIa, IIb (yellow) are functionally perfect flowers with functional anthers and non-functional stigmas; IIIc (white) is functionally sterile flower with non-functional anthers and stigma.

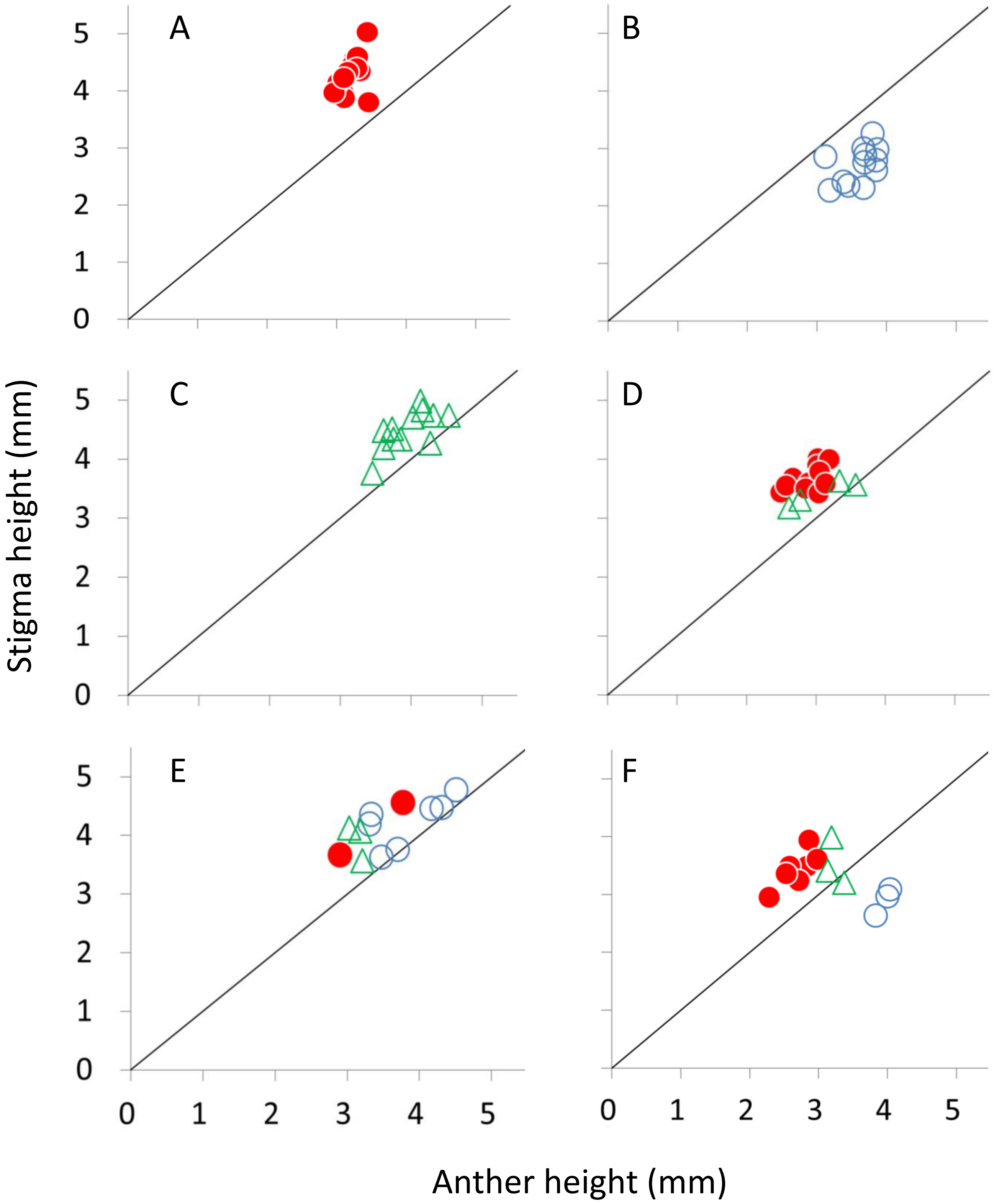


**FIG S2**

Anther/stigma height of female (closed circles), male (open circles), and perfect flowers (open triangles) in six plants (A-F) of *Psychotria manillensis* in the Sueyoshi population from Okinawa Island, Japan. A, a female plant only with female flowers; B, a male plant only with male flowers; C, a hermaphroditic plant only with perfect flowers; D, a monoecious plant with female and perfect flowers; E and F, monoecious plants with male, female and perfect flowers.

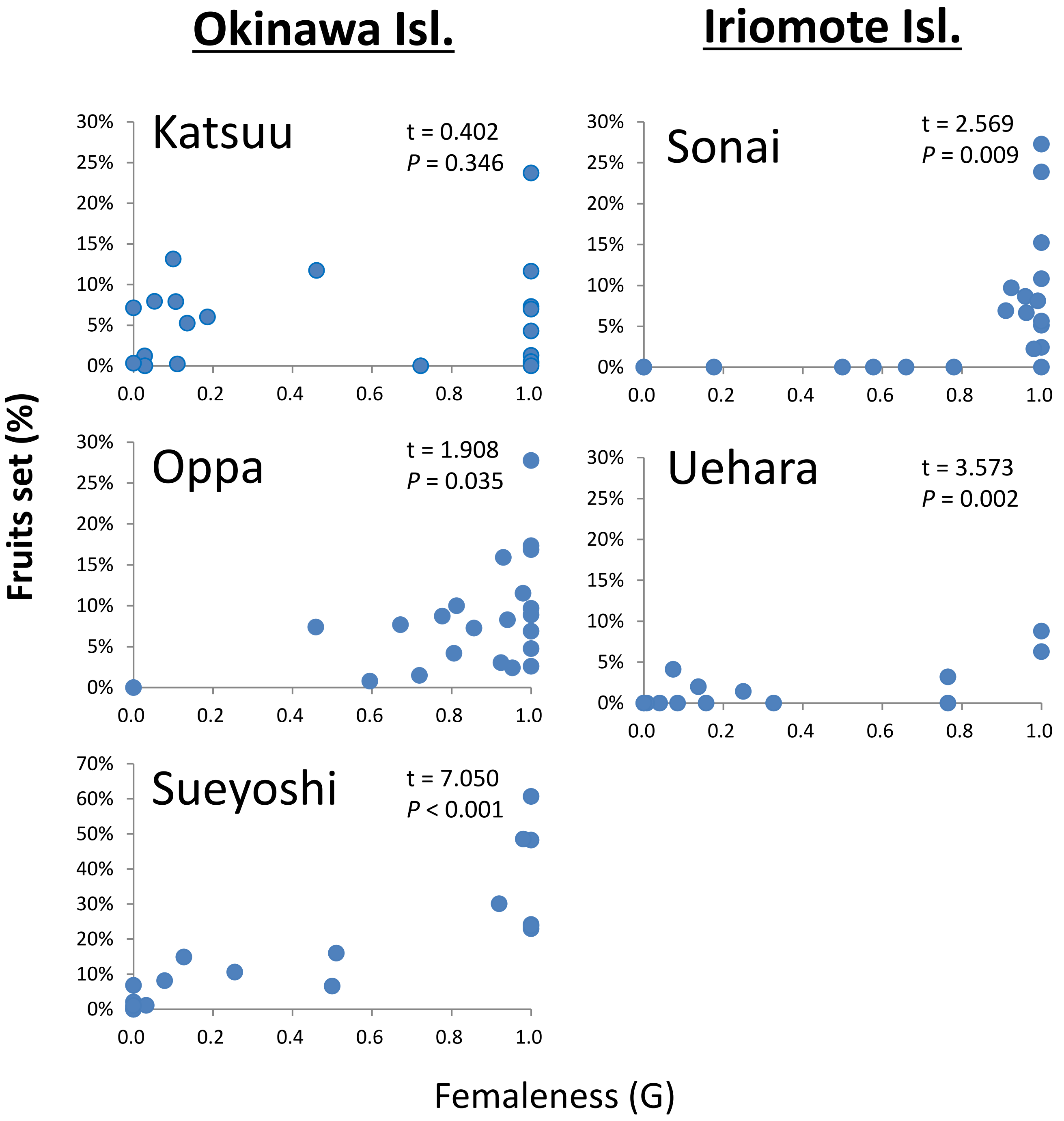


**Fig S3.**

Fruit set and femaleness (G) of *Psychotria manillensis* from the five populations on Okinawa and Iriomote Islands, Japan in 2012. Closed circles in the graphs represent individuals (20 from each population), straight lines are approximation lines, and t- and *P*-values are after Pearson's correlation test, assuming the correlation is greater than zero.

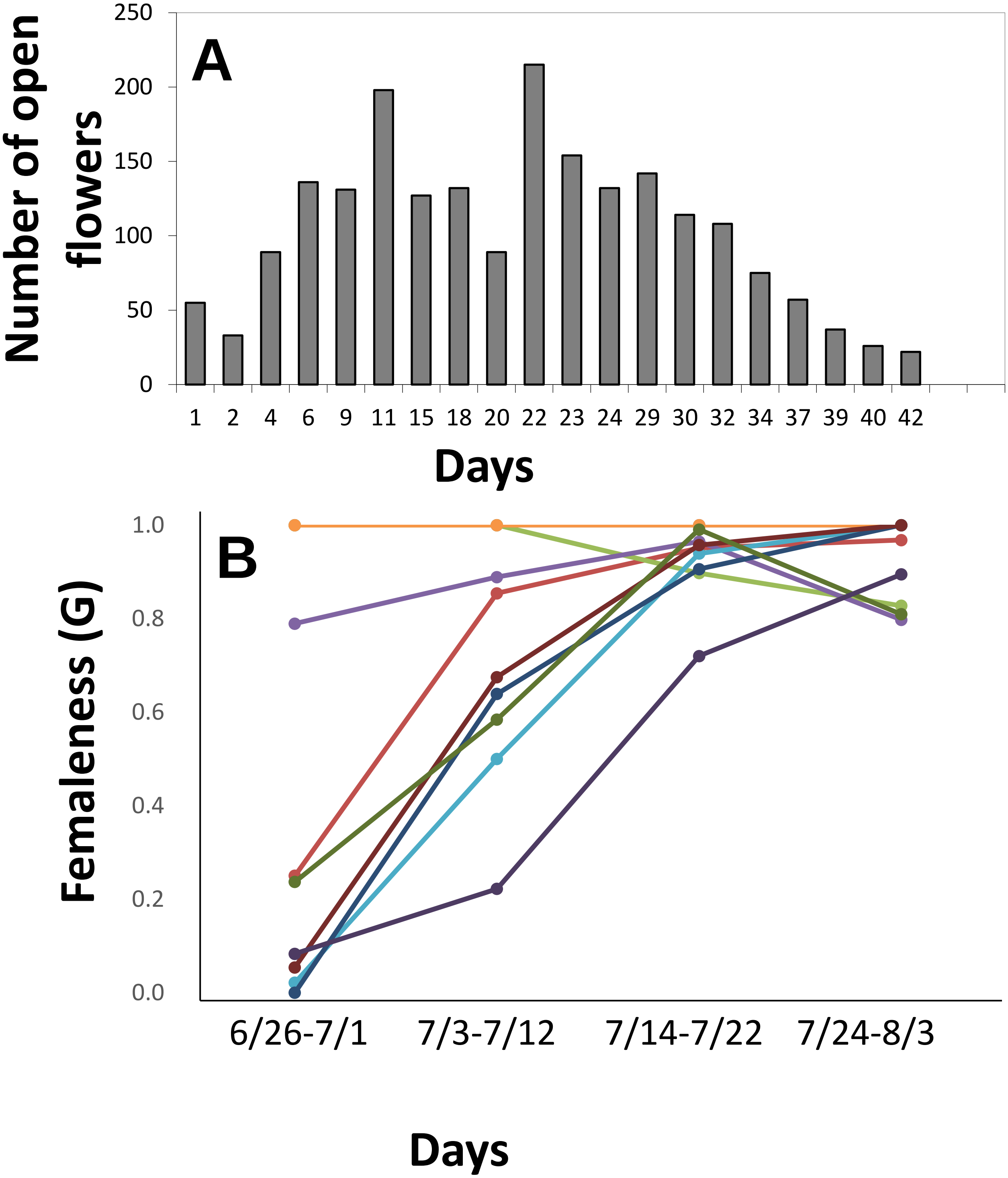


**FIG S4**

Flowering phenology (A) and relative femaleness (B) during the flowering period in 10 plants of *Psychotria manillensis* in the Oppa population from Okinawa Island, Japan in 2012. The number and sexual type of open flowers of 13 inflorescences on 10 individuals (130 inflorescences in total) every 2 d from June 26 to August 3, 2012. A: Number of open flowers showed the sum of all individuals which checked. B: Each line represents relative femaleness of each plant in each time span. Relative femaleness (G) in four time spans was calculated as follows: $G = (F + P/2)/(F + M + P)$, where F, P and M are the number of female, perfect and male flowers per plant, respectively. Two plants stay female throughout the seasons, thus those two lines are totally overlapped.
