## Supplemental table S1 for "Polygamous breeding system identified in the distylous genus *Psychotria*: *P. manillensis* in the Ryukyu archipelago, Japan"

Table S1. Fruit set of *Psychotria manillensis* in five populations of Okinawa and Iriomote Islands, Japan.

| | Population | N | *N* | n | Fruit set (%) | Percentage of fruiting plants |
|---|---|---|---|---|---|---|
| **Open pollination** | | | | | | |
| **2011** | Okinawa Isl. | | | | | |
| | Sueyoshi | 20 | 60 | 3138 | 12.5 | 100 |
| | Katsuu | 17 | 51 | 1828 | 21.9 | 100 |
| | Oppa | 33 | 99 | 4719 | 19.8 | 88.0 |
| **2012** | Okinawa Isl. | | | | | |
| | Sueyoshi | 20 | 60 | 4066 | 16.2 | 95.0 |
| | Katsuu | 23 | 61 | 6418 | 5.1 | 95.2 |
| | Oppa | 23 | 69 | 3896 | 8.2 | 95.7 |
| | Iriomote Isl. | | | | | |
| | Sonai | 20 | 58 | 2940 | 6.6 | 70.0 |
| | Uehara | 14 | 42 | 2135 | 1.8 | 64.3 |
| **2013** | Okinawa Isl. | | | | | |
| | Sueyoshi | 23 | 69 | 5064 | 6.9 | 100.0 |
| | Katsuu | 25 | 75 | 9061 | 12.0 | 100.0 |
| | Oppa | 22 | 66 | 6370 | 4.6 | 100.0 |
| | Iriomote Isl. | | | | | |
| | Sonai | 21 | 63 | 3818 | 3.3 | 95.2 |
| | Uehara | 13 | 39 | 1998 | 2.2 | 76.9 |
| **Bagging experiment** | | | | | | |
| **2012** | Okinawa Isl. | | | | | |
| | Sueyoshi | 10 | 10 | 444 | 0.68 | - |
| | Katsuu | 11 | 11 | 663 | 0.75 | - |
| | Oppa | 18 | 18 | 642 | 1.56 | |

N, number of plants

*N*, number of inflorescence

n, number of flowers
