## Supplemental table S2 for "Polygamous breeding system identified in the distylous genus *Psychotria*: *P. manillensis* in the Ryukyu archipelago, Japan"

Table S2. AIC of GLMM (logit-link, binomial distribution) and results of maximum likelihood test (chi-square) between models.

| | Models | Responsible variable | Fixed variable | Random effect | AIC | Maximum likelihood test between models | |
|---|---|---|---|---|---|---|---|
| | | | | | | Chisq | P value |
| Fruit set after cross pollination | | | | | | | |
| | model.0 | fruit set | none | plant id | 249 | 102.07 | <0.0001 |
| | model.1 | fruit set | stigma types | plant id | 151 | | |
| Fruit set after self- and cross polination | | | | | | | |
| | model.0 | fruit set | none | plant id + stigma type | 257 | 32.21 | <0.0001 |
| | model.1 | fruit set | treatment (cross/self) | plant id + stigma type | 227 | | |
| Fruit set under open pollination | | | | | | | |
| | model.0 | fruit set | none | plant id | 6953 | | |
| | model.1 | fruit set | year | plant id | 6809 | | |
| | model.2 | fruit set | population | plant id | 6914 | | |
| | model.3 | fruit set | year + population | plant id | 6780 | | |
| Fruit set after bagging and open pollination in 2012 | | | | | | | |
| | model.0 | fruit set | none | plant id / population | 2113 | 73.74 | <0.0001 |
| | model.1 | fruit set | treatment (bag/open) | plant id / population | 2041 | | |
