## Supplemental table S3 for "Polygamous breeding system identified in the distylous genus *Psychotria*: *P. manillensis* in the Ryukyu archipelago, Japan"

Table S3. Flower visitors of *Psychotria manillensis* on Okinawa and Iriomote Islands, the Ryukyu Islands, Japan, in 2011-2013.

| | | Population | | | | |
|---|---|---|---|---|---|---|
| | | Okinawa | | | Iriomote | |
| Family | Species | Kt | Op | Su | Ue | So |
| **HYMENOPTERA** | | | | | | |
| Apidae | *Amegilla dulcifera* | | 1 | 1 | | |
| | *A. florea* | | | | | 5 |
| | *Apis mellifera* | 2 | 2 | | | |
| Halictidae | *Lasioglossum* sp. | 1 | 2 | 4 | | |
| Icheumonidae | sp. | | 1 | | | |
| Scoliidae | *Megacampsomeris prismatica* | | | 1 | | |
| Vespidae | *Anterhynchium flavomarginatum* | 1 | | | | |
| | *Pararrhynchium ishigakiense* | | | | | 1 |
| | *Polistes formosanus* | | | | 22 | 16 |
| | *P. rothneyi* | 23 | 26 | 59 | 32 | 26 |
| | *Vespa affinis* | | | | 7 | 7 |
| | *V. analis* | 33 | 29 | 1 | 31 | 22 |
| | *V. ducalis* | | | | 7 | 5 |
| **DIPTERA** | | | | | | |
| Ephydridae | sp. | 1 | | | | |
| Calliphoridae | *Hemipyrellia* sp. | 2 | | | 1 | |
| | *Lucilia papuensis* | 1 | | | 1 | |
| | *L. porphyrina* | 12 | 22 | 10 | 9 | 4 |
| Muscidae | *Dichaetomyia flavipolis* | 3 | 7 | 5 | | |
| | *D. watasei* | 3 | 1 | | | |
| | *Stomoxys calcitrans* | 1 | | | | |
| Sarcophagidae | *Sarcophaga antilope* | | 1 | | | |
| | *S. kanakovi* | | 1 | | | |
| | sp. | 1 | | | | |
| Syrphidae | *Allobaccha nubilipennis* | 16 | 3 | 1 | 1 | |
| | sp. | | 2 | | | |
| Tachinidae | sp. | | 1 | | | |
| **LEPIDOPTERA** | | | | | | |
| Crambidae | sp. 1 | | 1 | | | |
| | sp. 2 | | 2 | | | |
| | sp. 3 | | 1 | | | |
| Lycaenidae | *Lampides boeticus* | 1 | | | | |
| Nymphalidae | *Athymaperius perius* | | 1 | | | |
| Papilionidae | *Papilio memnon* | 2 | | | | |
| | *Papilio okinawaensis* | 1 | | | | |
| | *Papilio polytes* | 4 | 1 | | | |
| Sphingidae | *Macroglossum pyrrhosticta* | | 2 | | | |
| **COLEOPTERA** | | | | | | |
| Chrysomelidae | *Altica cyanea* | 1 | | | | |
| **HEMIPTERA** | | | | | | |
| Largidae | *Physopelta cincticollis* | | 1 | | | |
| **ORTHOPTERA** | | | | | | |
| Gryllidae | *Cardiodactylus guttulus* | 1 | 1 | | 1 | 1 |
| | *Gryllodes sigillatus* | | | | | 2 |

Number of visits to an inflorescence during an 8-h observation are shown. Kt, Katsuu population; Op, Oppa population; Su, Sueyoshi population; Ue, Uehara population; So, Sonai population. Observations were performed on July 17 and 18, 2011 and on July 14 and 20, 2012 in the Katsuu population; from July 6 to September 18, 2011 and on July 14, 2012 in the Oppa population; on July 9 and 27, 2011, on June 19 and July 1, 2012 in the Sueyoshi population; and on June 2, 3, 15, 16, and 17, 2011 and on June 13, 2013 in the Sonai and Uehara populations.
